# Waning Immunity Against XBB.1.5 Following Bivalent mRNA Boosters

**DOI:** 10.1101/2023.01.22.525079

**Authors:** Ninaad Lasrado, Ai-ris Y. Collier, Jessica Miller, Nicole P. Hachmann, Jinyan Liu, Michaela Sciacca, Cindy Wu, Trisha Anand, Esther A. Bondzie, Jana L. Fisher, Camille R. Mazurek, Robert C. Patio, Olivia Powers, Stefanie L. Rodrigues, Marjorie Rowe, Nehalee Surve, Darren M. Ty, Bette Korber, Dan H. Barouch

**Affiliations:** Beth Israel Deaconess Medical Center, Boston, MA, USA; Los Alamos National Laboratory, Los Alamos, New Mexico, USA

## Abstract

The SARS-CoV-2 Omicron variant has continued to evolve. XBB is a recombinant between two BA.2 sublineages, XBB.1 includes the G252V mutation, and XBB.1.5 includes the G252V and F486P mutations. XBB.1.5 has rapidly increased in frequency and has become the dominant virus in New England. The bivalent mRNA vaccine boosters have been shown to increase neutralizing antibody (NAb) titers to multiple variants, but the durability of these responses remains to be determined. We assessed humoral and cellular immune responses in 30 participants who received the bivalent mRNA boosters and performed assays at baseline prior to boosting, at week 3 after boosting, and at month 3 after boosting. Our data demonstrate that XBB.1.5 substantially escapes NAb responses but not T cell responses after bivalent mRNA boosting. NAb titers to XBB.1 and XBB.1.5 were similar, suggesting that the F486P mutation confers greater transmissibility but not increased immune escape. By month 3, NAb titers to XBB.1 and XBB.1.5 declined essentially to baseline levels prior to boosting, while NAb titers to other variants declined less strikingly.

---

The SARS-CoV-2 Omicron variant has continued to evolve. XBB is a recombinant between two BA.2 sublineages, XBB.1 includes the G252V mutation, and XBB.1.5 includes the G252V and F486P mutations (**Fig. 1A**). XBB.1.5 has rapidly increased in frequency and has become the dominant virus in New England (**Fig. S1**). The bivalent mRNA vaccine boosters have been shown to increase neutralizing antibody (NAb) titers to multiple variants^1-4^, but the durability of these responses remains to be determined.

**Figure 1.**
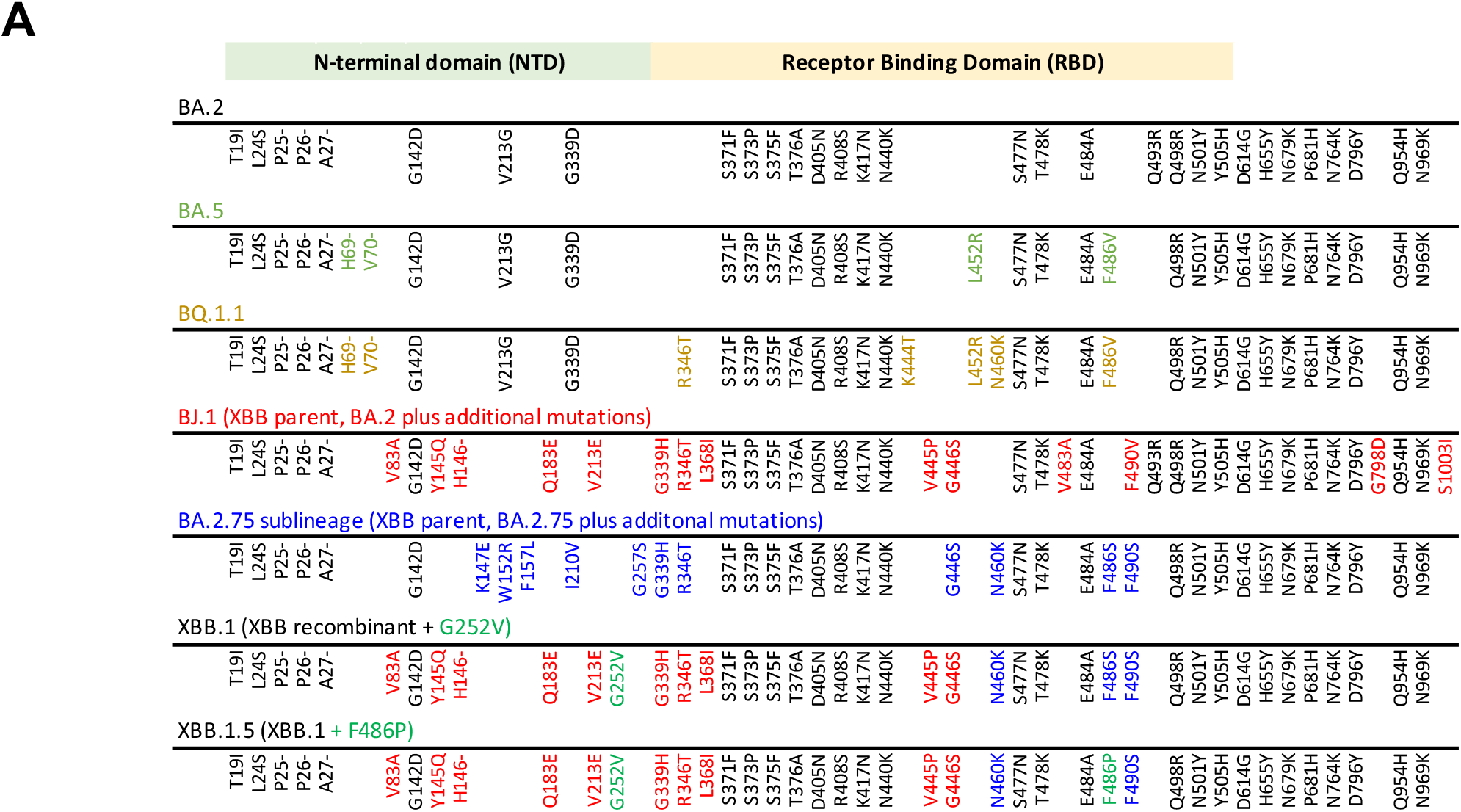
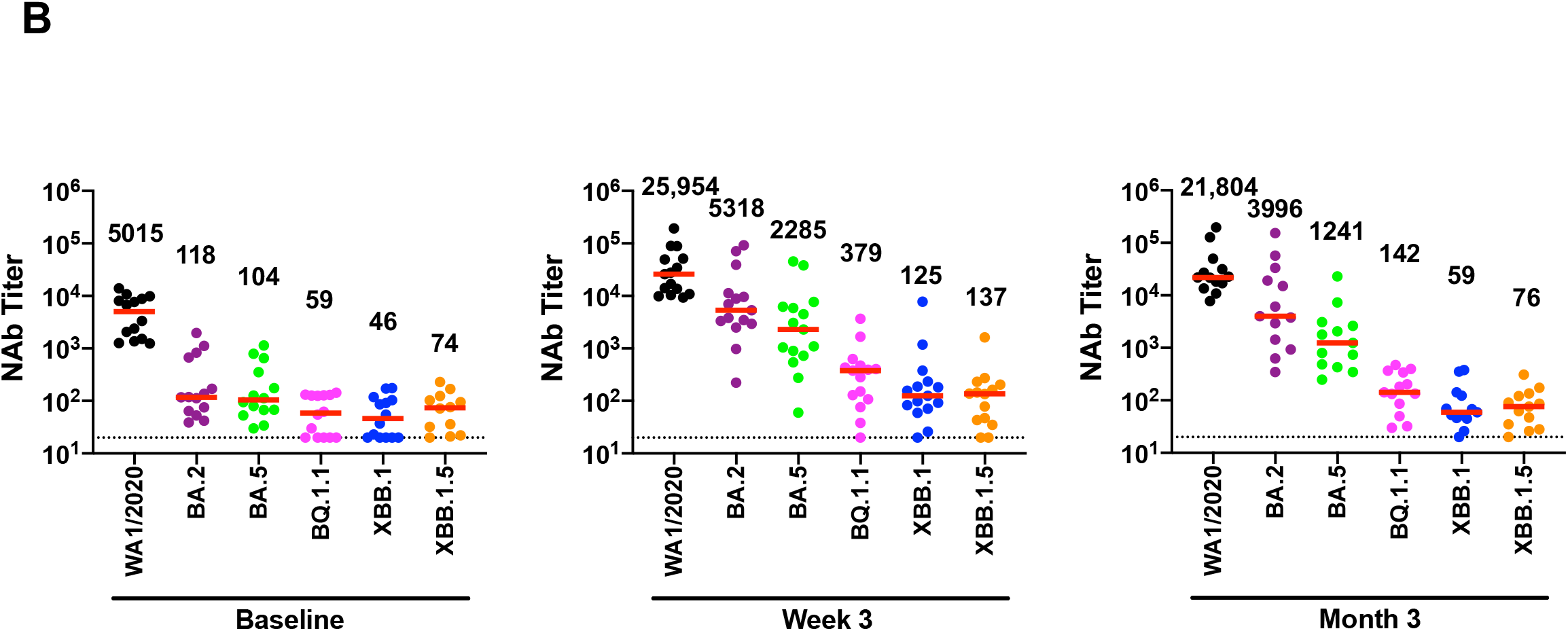
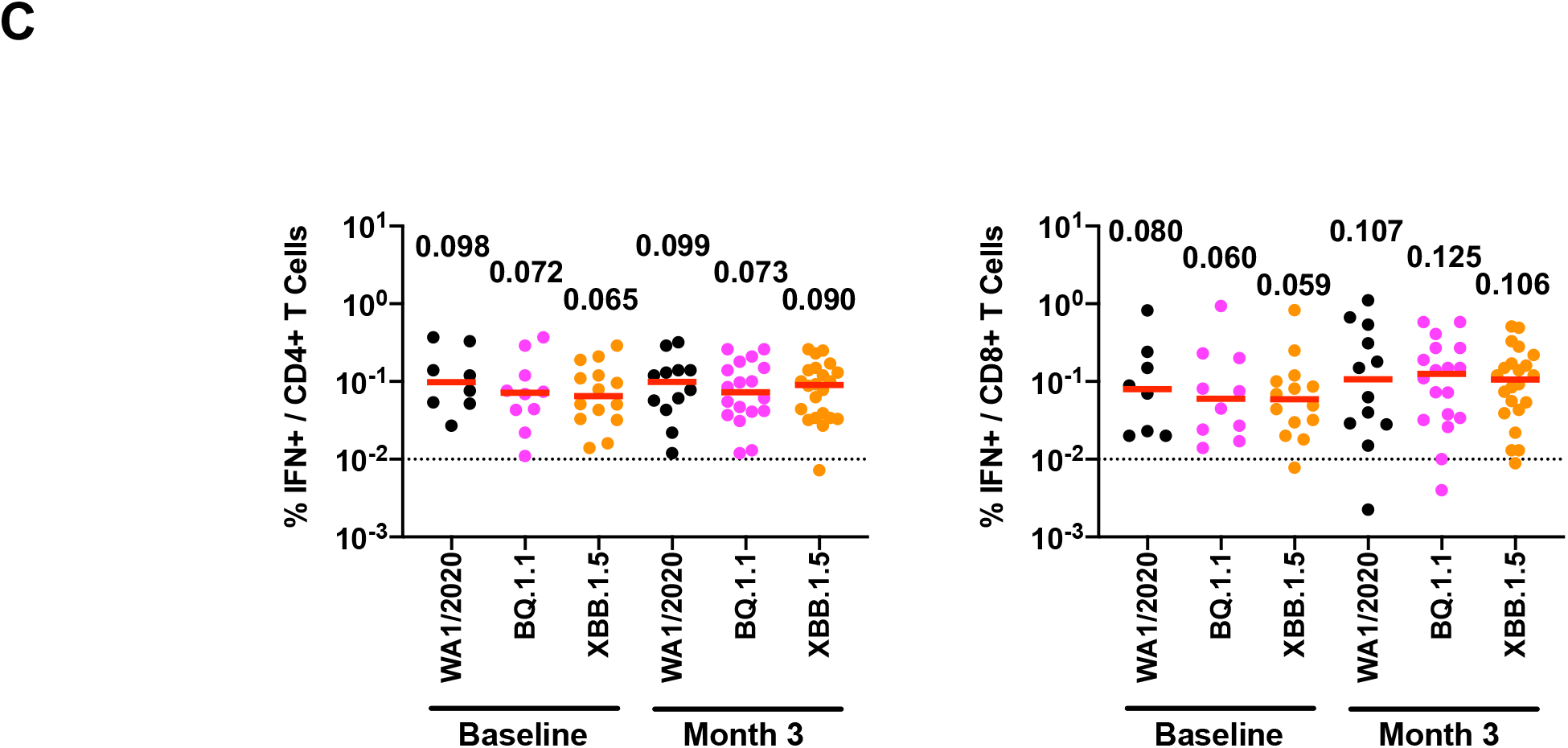
Humoral and cellular immune responses to SARS-CoV-2 Omicron variants. **A**. Spike sequences for BA.2, BA.5, BQ.1.1, XBB parental sequences BJ.1 (BA.2.10.1) and a sublineage of BA.2.75 (BM.1.1.1), XBB.1, and XBB.1.5 are depicted. Mutations compared with the ancestral WA1/2020 Spike are shown in black, and additional mutations relative to BA.2 are highlighted in colors corresponding to individual variants. For the XBB.1 and XBB.1.5 sequences, black reflects mutations from BA.2, red reflects additional mutations from BJ.1, blue reflects additional mutations from BA.2.75, and green reflects new mutations. NTD, N-terminal domain; RBD, receptor binding domain. **B**. Neutralizing antibody (NAb) titers against the WA1/2020, BA.2, BA.5, BQ.1.1, XBB.1, and XBB.1.5 variants by luciferase-based pseudovirus neutralization assays at baseline prior to boosting, at week 3 after boosting, and at month 3 after boosting in nucleocapsid seronegative participants. **C**. Spike-specific, IFN-γ CD4+ and CD8+ T cell responses to pooled WA1/2020, BQ.1.1, and XBB.1.5 peptides by intracellular cytokine staining assays at baseline prior to boosting and at month 3 after boosting. Dotted lines reflect limits of quantitation. Medians (red bars) are depicted and shown numerically.

We assessed humoral and cellular immune responses in 30 participants who received the bivalent mRNA boosters and performed assays at baseline prior to boosting, at week 3 after boosting, and at month 3 after boosting (**Table S1**). By month 3, 43% of participants had a known COVID-19 infection, although we speculate that this represents an underestimate of the true rate of infection. At baseline, median NAb titers to WA1/2020, BA.2, BA.5, BQ.1.1, XBB.1, and XBB.1.5 were 5015, 118, 104, 59, 46, and 74, respectively, in nucleocapsid seronegative participants (**Fig. 1B**). At week 3, median NAb titers to WA1/2020, BA.2, BA.5, BQ.1.1, XBB.1, and XBB.1.5 were 25,954, 5318, 2285, 379, 125, and 137, respectively (**Fig. 1B**). At month 3, median NAb titers to WA1/2020, BA.2, BA.5, BQ.1.1, XBB.1, and XBB.1.5 were 21,804, 3996, 1241, 142, 59, and 76, reflecting 1.2-, 1.3-, 1.8-, 2.7-, 2.1-, and 1.8-fold declines from week 3, respectively (**Fig. 1B**).

Spike-specific T cell responses were assessed by intracellular cytokine staining assays. Median CD4+ T cell responses to WA1/2020, BQ.1.1, and XBB.1.5 were 0.098%, 0.072%, and 0.065% at baseline and 0.099%, 0.073%, and 0.090% at month 3, respectively (**Fig. 1C**). Median CD8+ T cell responses to WA1/2020, BQ.1.1, and XBB.1.5 were 0.080%, 0.060%, and 0.059% at baseline and 0.107%, 0.125%, and 0.106% at month 3, respectively (**Fig. 1C**).

Our data demonstrate that XBB.1.5 substantially escapes NAb responses but not T cell responses after bivalent mRNA boosting. NAb titers to XBB.1 and XBB.1.5 were similar, suggesting that the F486P mutation confers greater transmissibility but not increased immune escape. By month 3, NAb titers to XBB.1 and XBB.1.5 declined essentially to baseline levels prior to boosting, while NAb titers to other variants declined less strikingly. The combination of low magnitude and rapidly waning NAb titers to XBB.1.5 will likely reduce the efficacy of the bivalent mRNA boosters^5^, but cross-reactive T cell responses, which were present prior to boosting, may continue to provide protection against severe disease.

---

**Table S1.** Study population

|  | **Bivalent mRNA Booster**<br>N=30 |
| --- | --- |
| **Age** (years), median (range) | 42 (24-77) |
| **Sex at birth**, Female | 21 (70) |
| **Race** |  |
| White | 27 (90) |
| Asian | 3 (10) |
| Black | 0 |
| More than one race | 0 |
| **Ethnicity** |  |
| Hispanic or Latino | 0 |
| Non-Hispanic | 30 (100) |
| **Medical condition** |  |
| Obesity (BMI $\geq 30$ kg/m<sup>2</sup>) | 6 (20) |
| Hypertension | 3 (10) |
| Diabetes | 1 (3) |
| Pregnant | 1 (3) |
| Asthma | 2 (7) |
| Living with HIV | 2 (7) |
| **Most recent COVID-19 vaccine** |  |
| Pfizer bivalent booster | 11 (37) |
| Moderna bivalent booster | 19 (63) |
| **Prior COVID-19 vaccines** |  |
| BNT (4 doses) | 3 (10) |
| BNT (3 doses) | 10 (33) |
| BNT (2 doses) / 1273 | 2 (7) |
| BNT (2 doses) / Ad26 / BNT | 1 (3) |
| BNT (2 doses) / Ad26 / 1273 | 1 (3) |
| 1273 (4 doses) | 2 (7) |
| 1273 (3 doses) | 9 (30) |
| Ad26 / BNT (2 dose) / 1273 | 1 (3) |
| Ad26 / 1273 (1 dose) | 1 (3) |
| **Interval between last two vaccine doses (days)** | 326 (250-364) |
| **Days from bivalent vaccine dose to initial sampling** | 21 (16-24) |
| **Days from bivalent vaccine dose to 3 mo sampling** | 92 (86-96) |
| **Known COVID-19 positive\*** | 13 (43) |

BNT=BNT162b2; 1273=mRNA-1273; Ad26=Ad26.COV2.S
Data displayed as median (range or interquartile range, IQR) and n (%); BMI, body mass index; pregnant designation reflects time of last vaccine dose and/or time of sampling.
\* Two participants had infections between day 7-40 after the bivalent booster.

## Supplementary Figure Legend

**Figure S1.**
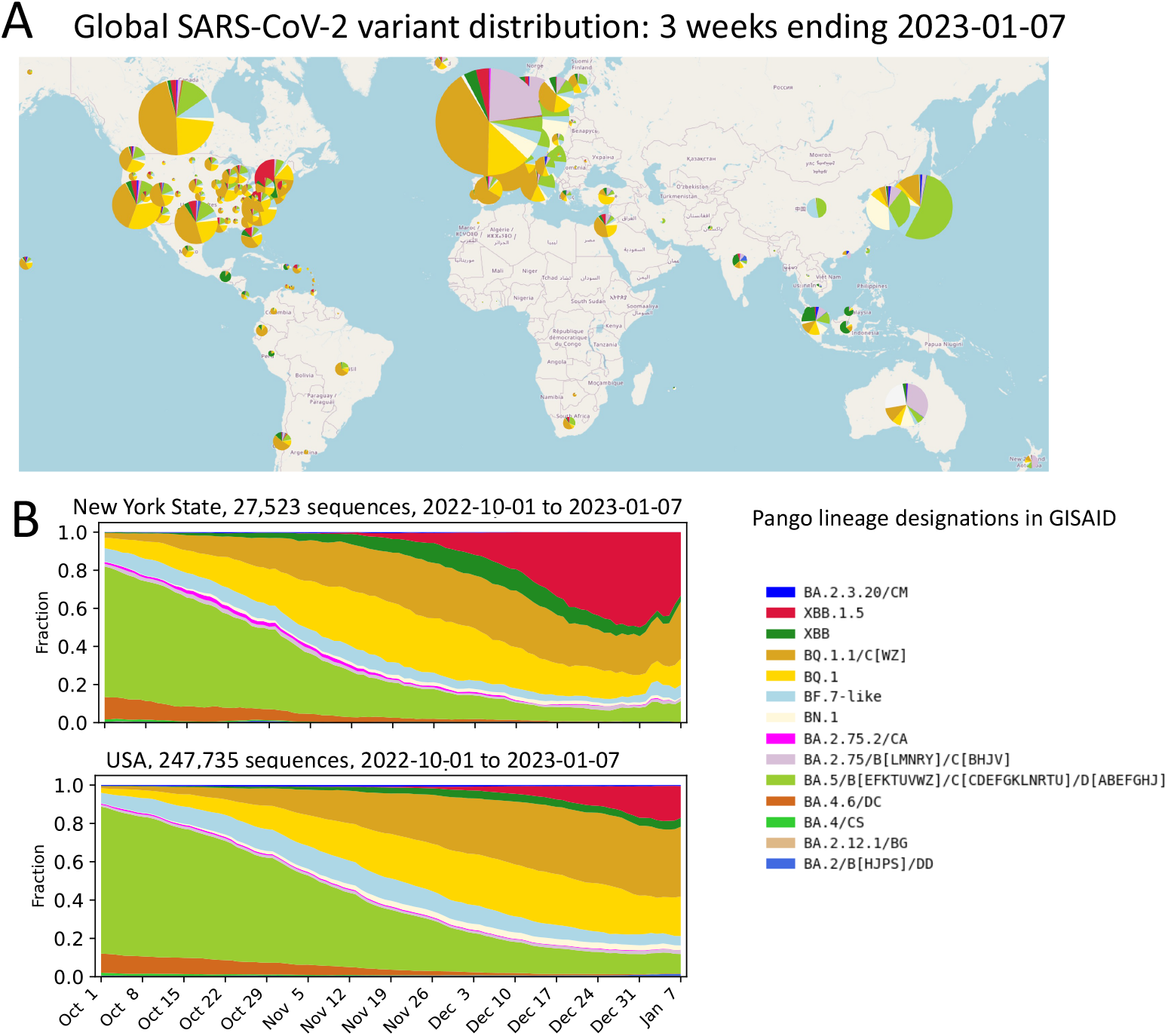
Increasing prevalence of the XBB.1.5 variant. **A**. Global frequencies of circulating Pango lineages. Pango lineage designations assigned to GISAID sequences, country of origin, and date of sampling were used to generate these figures. Sequences available in GISAID as of 2023-01-21 were included, but analyses were restricted to samples obtained through 2023-01-07, as the natural delay between sampling and availability in GISAID makes very recent data sparce and prone to sampling bias. Data visualizations were created at cov.lanl.gov. XBB.1.5, shown in red, has rapidly increased in frequency, particularly in the Northeastern United States. The earliest XBB.1.5 variant was sampled in New York on 2022-10-22. As of 2023-01-21, XBB.1.5 is established (sampled 10 times or more) in 24 countries and is increasing in sampling frequency everywhere it has been introduced. Note, however, that during this same period other BA.2.75 lineages including BN.1, other XBB sublineages, and distinctive BA.5 sub-lineages have also been gaining prevalence in other parts of the world, and the current global variant picture is complex. **B**. The rapid increase in sampling frequency of XBB.1.5 in New York and throughout the United States between 2022-10-01 and 2023-01-07. Current models based on NOWCAST estimates from the Centers for Disease Control and Prevention (CDC) variant tracking tools suggest that XBB.1.5 may have reached 49% (38-61%) in the USA and 87% (83-90%) in the region that includes New York. The actual frequencies based on GISAID data is shown here, and these XBB.1.5 frequencies are slightly lower than the CDC estimates.

## Supplementary Methods

### Study Population

A specimen biorepository at Beth Israel Deaconess Medical Center (BIDMC) obtained samples from individuals who received SARS-CoV-2 vaccines as well as bivalent mRNA boosters. The BIDMC institutional review board approved this study (2020P000361). All participants provided informed consent. This study included 30 individuals who received the bivalent mRNA boosters (Pfizer-BioNTech, Moderna). Participants were excluded from the immunologic assays if they had a history of SARS-CoV-2 infection or a positive nucleocapsid (N) serology by electrochemiluminescence assays (ECLA) or if they received immunosuppressive medications.

### Pseudovirus Neutralizing Antibody Assay

Neutralizing antibody (NAb) titers against SARS-CoV-2 variants utilized pseudoviruses expressing a luciferase reporter gene. In brief, the packaging construct psPAX2 (AIDS Resource and Reagent Program), luciferase reporter plasmid pLenti-CMV Puro-Luc (Addgene), and Spike protein expressing pcDNA3.1-SARS-CoV-2 SΔCT were co-transfected into HEK293T cells (ATCC CRL_3216) with lipofectamine 2000 (ThermoFisher Scientific). Pseudoviruses of SARS-CoV-2 variants were generated using the Spike protein from WA1/2020 (Wuhan/WIV04/2019, GISAID accession ID: EPI_ISL_402124), Omicron BA.2 (GISAID ID: EPI_ISL_6795834.2), BA.5 (GISAID ID: EPI_ISL_12268495.2), BQ.1.1 (GISAID ID: EPI_ISL_14752457), XBB.1 (GISAID ID: EPI_ISL_15232105), and XBB.1.5 (GISAID ID: EPI_ISL_16418320). The supernatants containing the pseudotype viruses were collected 48h after transfection, and pseudotype viruses were purified by filtration with 0.45-μm filter. To determine NAb titers in human serum, HEK293T-hACE2 cells were seeded in 96-well tissue culture plates at a density of 2 × 10^4^ cells per well overnight. Three-fold serial dilutions of heat-inactivated serum samples were prepared and mixed with 60 μl of pseudovirus. The mixture was incubated at 37 °C for 1 h before adding to HEK293T-hACE2 cells. After 48 h, cells were lysed in Steady-Glo Luciferase Assay (Promega) according to the manufacturer’s instructions. SARS-CoV-2 neutralization titers were defined as the sample dilution at which a 50% reduction (NT50) in relative light units was observed relative to the average of the virus control wells.

### Intracellular cytokine staining (ICS) assay

CD4+ and CD8+ T cell responses were quantitated by pooled peptide-stimulated intracellular cytokine staining (ICS) assays. Peptide pools contained 15 amino acid peptides overlapping by 11 amino acids spanning the SARS-CoV-2 WA1/2020, BQ.1.1, or XBB.1.5 Spike proteins (21st Century Biochemicals). 10^6^ peripheral blood mononuclear cells were re-suspended in 100 μL of R10 media supplemented with CD49d monoclonal antibody (1 μg/mL) and CD28 monoclonal antibody (1 μg/mL). Each sample was assessed with mock (100 μL of R10 plus 0.5% DMSO; background control), peptides (2 μg/mL), and/or 10 pg/mL phorbol myristate acetate (PMA) and 1 μg/mL ionomycin (Sigma-Aldrich) (100μL; positive control) and incubated at 37°C for 1 h. After incubation, 0.25 μL of GolgiStop and 0.25 μL of GolgiPlug in 50 μL of R10 was added to each well and incubated at 37°C for 8 h and then held at 4°C overnight. The next day, the cells were washed twice with DPBS, stained with aqua live/dead dye for 10 mins and then stained with predetermined titers of monoclonal antibodies against CD279 (clone EH12.1, BB700), CD4 (clone L200, BV711), CD27 (clone M-T271, BUV563), CD8 (clone SK1, BUV805), CD45RA (clone 5H9, APC H7) for 30 min. Cells were then washed twice with 2% FBS/DPBS buffer and incubated for 15 min with 200 μL of BD CytoFix/CytoPerm Fixation/Permeabilization solution. Cells were washed twice with 1X Perm Wash buffer (BD Perm/WashTM Buffer 10X in the CytoFix/CytoPerm Fixation/ Permeabilization kit diluted with MilliQ water and passed through 0.22μm filter) and stained intracellularly with monoclonal antibodies against Ki67 (clone B56, BB515), IL21 (clone 3A3-N2.1, PE), CD69 (clone TP1.55.3, ECD), IL10 (clone JES3-9D7, PE CY7), IL13 (clone JES10-5A2, BV421), IL4 (clone MP4-25D2, BV605), TNF-α (clone Mab11, BV650), IL17 (clone N49-653, BV750), IFN-γ (clone B27; BUV395), IL2 (clone MQ1-17H12, BUV737), IL6 (clone MQ2-13A5, APC), and CD3 (clone SP34.2, Alexa 700) for 30 min. Cells were washed twice with 1X Perm Wash buffer and fixed with 250μL of freshly prepared 1.5% formaldehyde. Fixed cells were transferred to 96-well round bottom plate and analyzed by BD FACSymphony™ system. Data were analyzed using FlowJo v9.9.

### Data availability

All genome sequences and associated metadata in this dataset are published in GISAID’s EpiCoV database. The contributors of each individual sequence with details such as accession number, virus name, collection date, originating lab, and submitting lab and the list of authors can be found as detailed below.

GISAID data used in Figure S1A: GISAID Identifier: EPI_SET_230122qg, doi: 10.55876/gis8.230122qg, EPI_SET_230122qg is composed of 123,538 individual genome sequences. The collection dates range from 2022-12-17 to 2023-01-07. Data were collected in 78 countries and territories.

GISAID data used in Figure S1B: GISAID Identifier: EPI_SET_230122yz, doi: 10.55876/gis8.230122yz, EPI_SET_230122yz is composed of 234,576 individual genome sequences. The collection dates range from 2022-10-01 to 2023-01-07. Data were collected in 7 countries and territories.

We gratefully acknowledge all the people who contribute sequences and related data to GISAID, enabling a global portrait of emerging SARS-CoV-2 variants, and the staff at GISAID (https://gisaid.org) and at the Los Alamos Laboratory COVID-19 Viral Genome Analysis Pipeline (https://cov.lanl.gov/content/index) for providing the bioinformatic tools that enable rapid analyses.

